# Broadband backscattering confocal microscopy enables label-free 3D live cell nanoscale sensitive imaging

**DOI:** 10.64898/2026.02.02.703335

**Authors:** Mark F. Coughlan, Lei Zhang, Rebecca T. Perelman, Umar Khan, Xuejun Zhang, Paul K. Upputuri, Yuri N. Zakharov, Le Qiu, Lev T. Perelman

## Abstract

Fluorescence microscopy is a cornerstone of biological research. However, fluorescent labeling is challenging in live cells and is constrained by photobleaching and phototoxicity. Label-free methods allow cells to be studied in their native state, but most techniques have poor contrast, lack 3D capability, rely on complex optics, and fail to provide structural information. We present broadband backscattering confocal microscopy (BBCM), which employs a broadband supercontinuum laser and collects backscattered light in confocal geometry using a photomultiplier tube. Broadband illumination averages out size-dependent oscillations that confound monochromatic backscattering. This eliminates blind spots and intensity ambiguities, allowing all scatterers to be visible, with the signal increasing approximately linearly with scatterer size. BBCM is easy to retrofit to standard confocal microscopes, requires no specialized optics, and is straightforward for nonspecialists. It enables high-contrast, label-free 3D imaging of live cells with size sensitivity to subcellular structures without employing custom optics or complex data processing.

---

Optical microscopy provides the most direct method for probing the structure, function, and physiology of living systems. Numerous microscopy advancements over the past 400 years have fundamentally changed the way cells and molecules are examined^1^. Fluorescence microscopy remains the most widely used technique, mainly due to its high contrast and molecular specificity^2,3^. Combining fluorescence imaging with confocal configurations dramatically reduces out-of-focus light, further improving optical contrast and enabling diffraction-limited optical sectioning^4,5^. Despite the widespread utility of fluorescence imaging, its use in live cells is limited by illumination-induced phototoxicity, photobleaching, perturbations from exogenous labels, and the incompatibility of fixation-dependent labeling strategies^6,7^.

Given these drawbacks, there is a fundamental need for label-free approaches that can visualize live cells in their native unperturbed state^8-10^. The most accessible label-free modality is bright-field microscopy, which detects intensity variations in transmitted light. However, because cells are primarily phase objects, they have low intrinsic contrast and appear faint. Dark-field microscopy is perhaps the simplest contrast enhancing label-free approach^11,12^. The main principle is the collection of only object-scattered light, while eliminating the unscattered illumination light by means of a special illumination design. Dark-field microscopy is simple to retrofit, highly sensitive to subwavelength scatterers, and compatible with live specimens. However, it is limited to two-dimensional analysis and loses signal rapidly with depth or in strongly scattering media.

Phase-contrast microscopy, introduced by Zernike^13,14^, uses a phase plate to selectively interfere the object-scattered light with the unscattered illumination light. This yields high-contrast images of transparent specimens and remains a staple for monitoring cell proliferation and morphology^15^. However, halo artifacts can obscure structural boundaries, and axial sectioning is limited by the wide-field geometry.

Another widely adopted contrast-enhancing technique is differential interference contrast (DIC) microscopy^16,17^, which employs a polarized light beam separated into two laterally displaced, orthogonally polarized components that interfere after passing through the specimen. DIC provides excellent edge contrast and axial resolution. Nonetheless, image contrast depends on the orientation of optical gradients, and birefringent optics can hinder multimodal integration. As with dark-field and phase contrast, the measurement remains inherently two-dimensional.

Quantitative phase imaging (QPI)^18^, which includes techniques such as digital holographic microscopy^19^, diffraction phase microscopy^20^, and tomographic phase microscopy^21^, overcomes these limitations by measuring the absolute phase delay introduced by the specimen. This enables quantitative mapping of changes in optical path length, thereby facilitating accurate and dynamic visualization of cellular morphology in live specimens. While QPI has provided significant insights across biological and medical research^22,23^, it remains difficult for non-specialists because of the required interferometric optics, alignment tolerances, and phase-unwrapping algorithms. Furthermore, it provides rather limited structural information.

Thus, despite the broad range of label-free contrast-enhancing techniques available, there remains a critical need for methods that provide straightforward and accessible high-contrast three-dimensional structure-sensitive imaging of live cells. Here, we present broadband backscattering confocal microscopy (BBCM), a technique that collects broadband sample-scattered light in a confocal backscattering configuration. BBCM enables three-dimensional label-free imaging of live cells with high optical contrast and size sensitivity to subcellular compartments, providing a platform for long-term volumetric interrogation of cellular structures and dynamics.

## RESULTS

### Operating principle of BBCM

The BBCM setup is shown in Fig. 1a. It employs a standard point-scanning confocal microscope, with a supercontinuum laser source coupled through free space in addition to conventional monochromatic lasers. The supercontinuum source, a coherent white light laser that emits light in the 425 nm to 2000 nm range, complements traditional lasers by enabling broadband spectral acquisition within the same system. Infrared light is removed using an external absorption filter, and the entire visible band is transmitted through the microscope via a beamsplitter mounted in one of the slots on the scan-head dichroic wheel. The visible light is then focused into a tight focal volume using a plan-apochromat objective, as illustrated in Fig. 1b. When a scatterer enters the focal volume of the white light beam (Fig. 1b inset), light is scattered in many directions, with most of the light scattered in the forward direction. Crucially, a portion of the backscattered light is collected by the objective, while out-of-focus backscattered light is rejected by the detection pinhole, thereby ensuring optical sectioning.

**Fig. 1.**
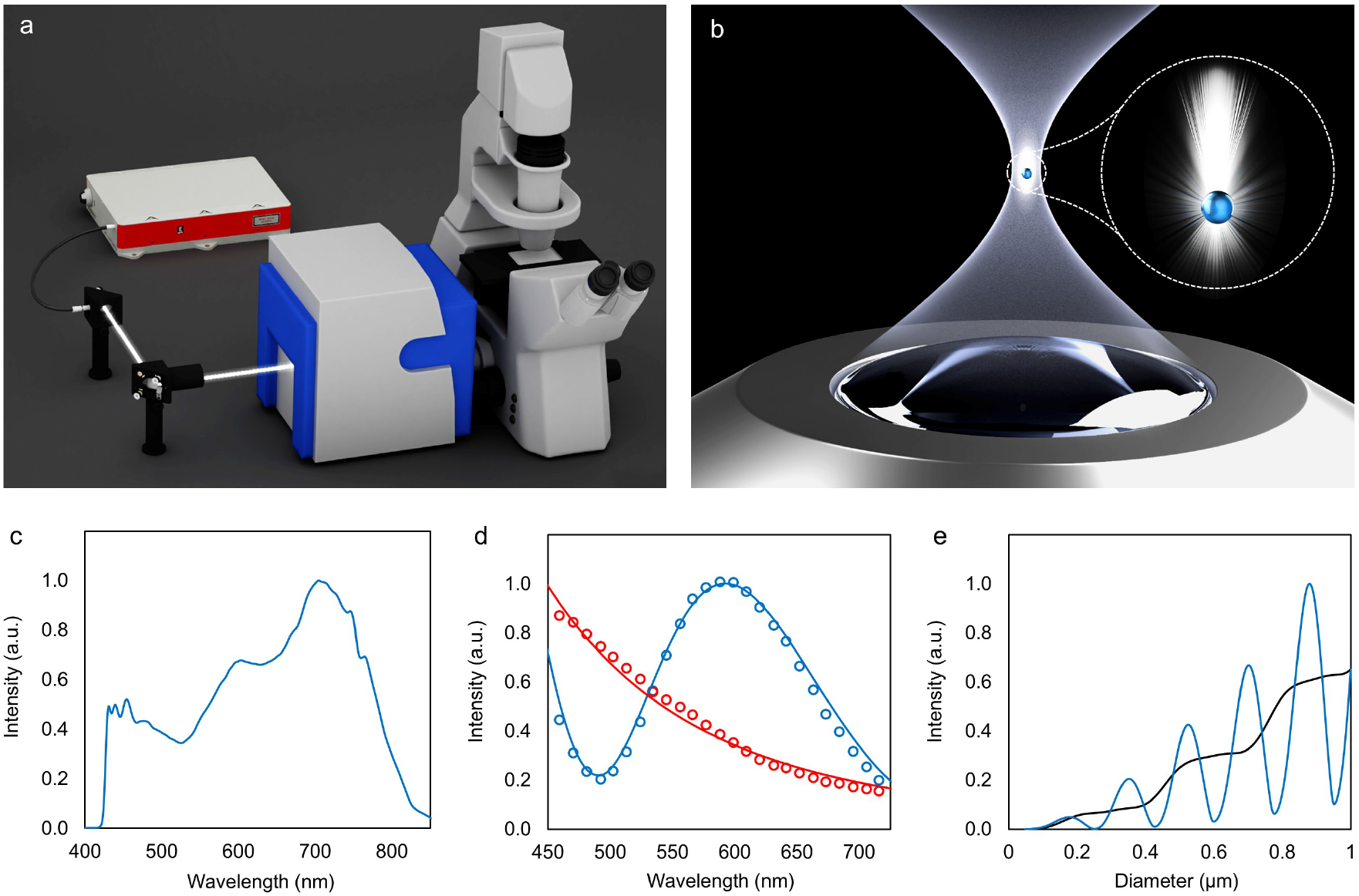
Broadband backscattering confocal microscopy (BBCM) setup and principle of operation. **a**, A supercontinuum white light source is coupled into the point scanning confocal microscope, with a beamsplitter mounted in one of the slots on the scan-head dichroic wheel. **b**, The plan-apochromat objective focuses the coherent white light into a small focal volume. The inset shows scattered light from a small spherical particle, with the backscattered light collected by the objective. **c**, The spectrum that emerges from the objective shows that light across the entire visible range is transmitted through the microscope. **d**, Light scattering spectra for 100 nm diameter polystyrene microspheres (red) and 350 nm diameter polystyrene microspheres (blue). Open circles show experimental data, while solid lines show model fits. **e**, Simulated scattering intensity for spherical particles of different diameters. The blue curve represents monochromatic illumination at 488 nm, while the black curve corresponds to broadband illumination spanning the 425 to 800 nm range.

The spectrum emerging from the objective is shown in Fig. 1c, demonstrating a continuous white-light output spanning 425 to 800 nm. While a monochromatic laser also produces scattered light, broadband illumination allows scatterers of many sizes to contribute to the signal, improving sensitivity across spatial scales. Wavelength-dependent scattering from subcellular structures of varying sizes has previously been exploited both in non-microscopy methods, such as light scattering spectroscopy (LSS)^24,25^, and in microscopy-based methods, such as confocal light absorption and scattering spectroscopic (CLASS) microscopy^26,27^. In the latter, tight beam focusing confines the scattering measurement to a well-defined confocal volume. While both LSS and CLASS microscopy rely on spectrally resolved measurements and light-scattering models, BBCM instead records the entire broadband signal with a single photomultiplier tube (PMT), eliminating the need for spectral processing and computational modeling. However, the physics of light scattering and confocal microscopy detection underlying CLASS microscopy provide a direct conceptual basis for understanding the operating principle of BBCM.

Figure 1d shows the experimental spectra acquired using CLASS microscopy for polystyrene microspheres 100 nm and 350 nm in diameter, and corresponding fits to a previously established light-scattering model^28^ (briefly described in the Supplementary Information). The excellent agreement between model fits and experimental data confirms that scattering within the confocal volume is accurately characterized, enabling reliable simulation of the total scattered light from spherical particles of varying diameters, as shown in Fig. 1e. This simulation represents the backscattered signal collected by a confocal microscope with a single PMT, as implemented in BBCM. Figure 1e shows scattering intensity from spherical particles as a function of diameter under monochromatic illumination at 488 nm and broadband illumination from a supercontinuum laser with the same incident power. The monochromatic illumination produces pronounced size-dependent oscillations in backscattering intensity, whereas the broadband illumination yields a near-linear increase with particle diameter. At certain sizes, the monochromatic illumination produces minimal scattering, rendering those particles effectively invisible.

Furthermore, scatterers of very different sizes can produce identical backscattering intensities, making monochromatic backscattering contrast inherently ambiguous in standard confocal systems. In contrast, broadband illumination produces measurable scattering for particles of all sizes, with intensity that scales approximately linearly with diameter, ensuring consistent visibility and size-resolved contrast. To further illustrate this, we modeled the backscattering intensity for several common laser wavelengths (405 nm, 458 nm, 514 nm, 543 nm, 594 nm, and 633 nm) as shown in Supplementary Fig. 1. Although the number and spacing of oscillations vary with wavelength, each wavelength has characteristic sizes that scatter little light and other sizes that give identical backscattering intensities, preventing backscattering intensity alone from distinguishing particle sizes.

In summary, BBCM employs a supercontinuum laser to focus coherent white light into a tight focal volume and collects the backscattered light from scatterers within that volume in a confocal configuration. The broadband nature of the illumination ensures that a wide range of scatterer sizes and types contribute to the signal. Importantly, in BBCM, backscattered signals increase approximately linearly with size, making scatterers both visible and readily distinguishable by backscattering intensity.

### High-contrast label-free cell measurements

Figure 2a shows an image of PANC-1 cells obtained with BBCM. The confocal configuration provides excellent optical sectioning, collecting only backscattered light from a thin optical slice within the cells. This enables label-free visualization of nuclei, cell boundaries, and intracellular organelles with high optical contrast. To verify that the observed structures corresponded to nuclei, co-registered fluorescence images were acquired from the same field of view (Fig. 2b). The merged images (Fig. 2c) show excellent spatial overlap, confirming that BBCM can be used for label-free imaging of cell nuclei. Figures 2d-f shows another example of BBCM, fluorescence, and merged images, respectively. The ability to visualize nuclei without labels is a major advantage, as nuclei are convenient and information-rich imaging targets in cell biology. Their consistent shape and well-defined positions make them valuable for segmenting individual cells and monitoring their movement or division over time^29,30^. Moreover, nuclear morphology can reflect key biological processes, including mitosis, differentiation, and cell death^31,32^.

**Fig. 2.**
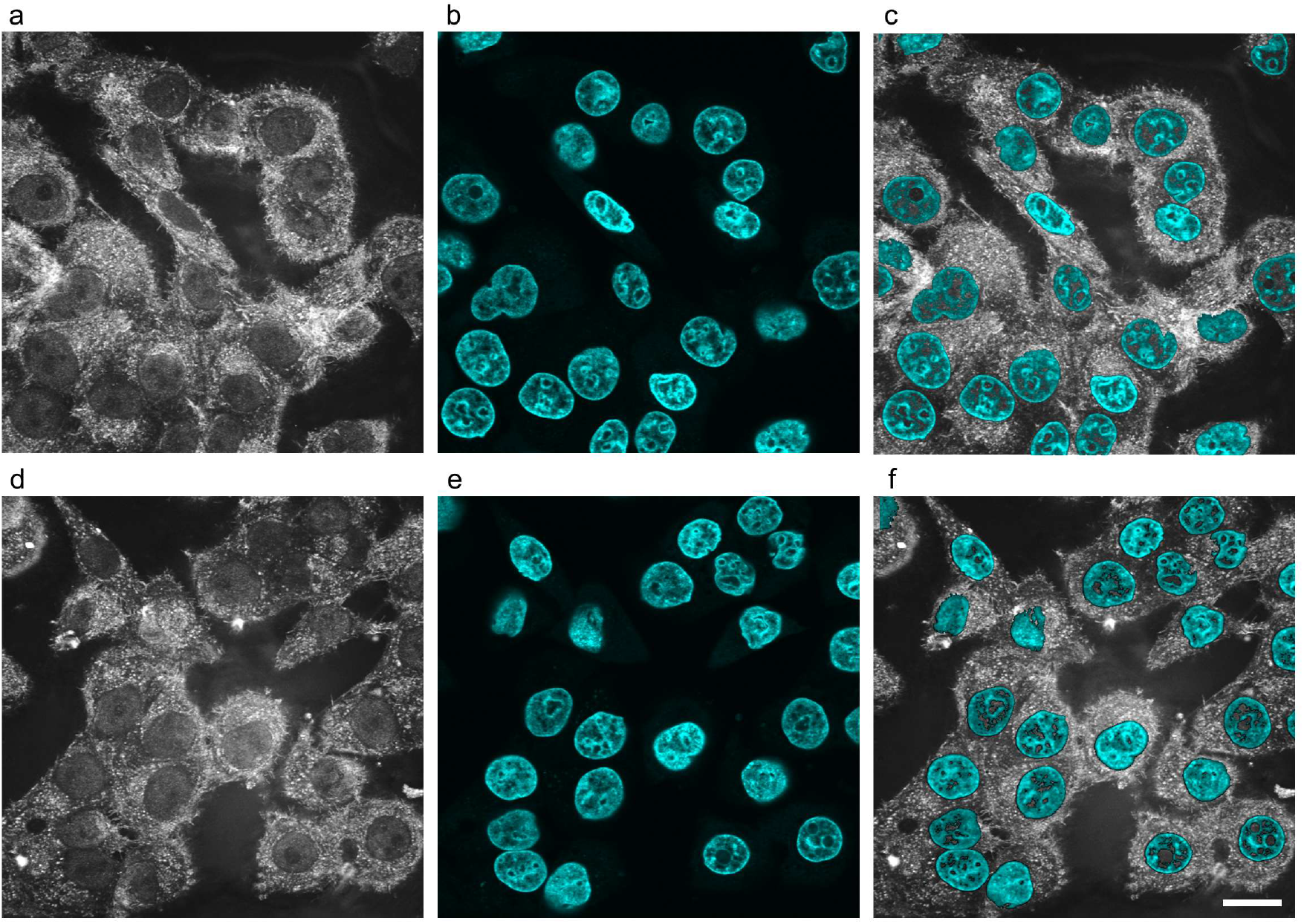
BBCM images of PANC-1 cells with co-registered fluorescence images of cell nuclei. **a**, BBCM image of PANC-1 cells showing clear cell and nuclear outlines. **b**, Fluorescence image from the same field of view as (a) showing DAPI-stained cell nuclei. **c**, Merged BBCM and fluorescence images showing strong spatial overlap in nuclear regions. **d**, Same as (a). **e**, Fluorescence image acquired from the same field of view as in (d). **f**, Merged BBCM and fluorescence images. Scale bar 20 µm.

To evaluate whether BBCM can also resolve cell outlines, we imaged HeLa cells double-stained for nuclei and cytoplasm. BBCM images are shown in Figs. 3a and 3d, and the corresponding pseudo-stained images are shown in Figs. 3b and 3e. To generate these pseudo-stains, a local entropy filter was first applied to the BBCM images to enhance structural detail. A red colormap with an offset factor was then used to visualize cytoplasmic features. A binary nuclear mask obtained from prior segmentation was used to isolate nuclear and non-nuclear regions. Intensities within the nuclear region were normalized and visualized with a cyan colormap. Additional details on the pseudo-staining procedure are provided in the *Methods* section. The corresponding fluorescence images (Figs. 3c and 3f) show strong agreement with the pseudo-stained BBCM images (Figs. 3b and 3e), confirming that both nuclear and cytoplasmic structures are accurately captured.

**Fig. 3.**
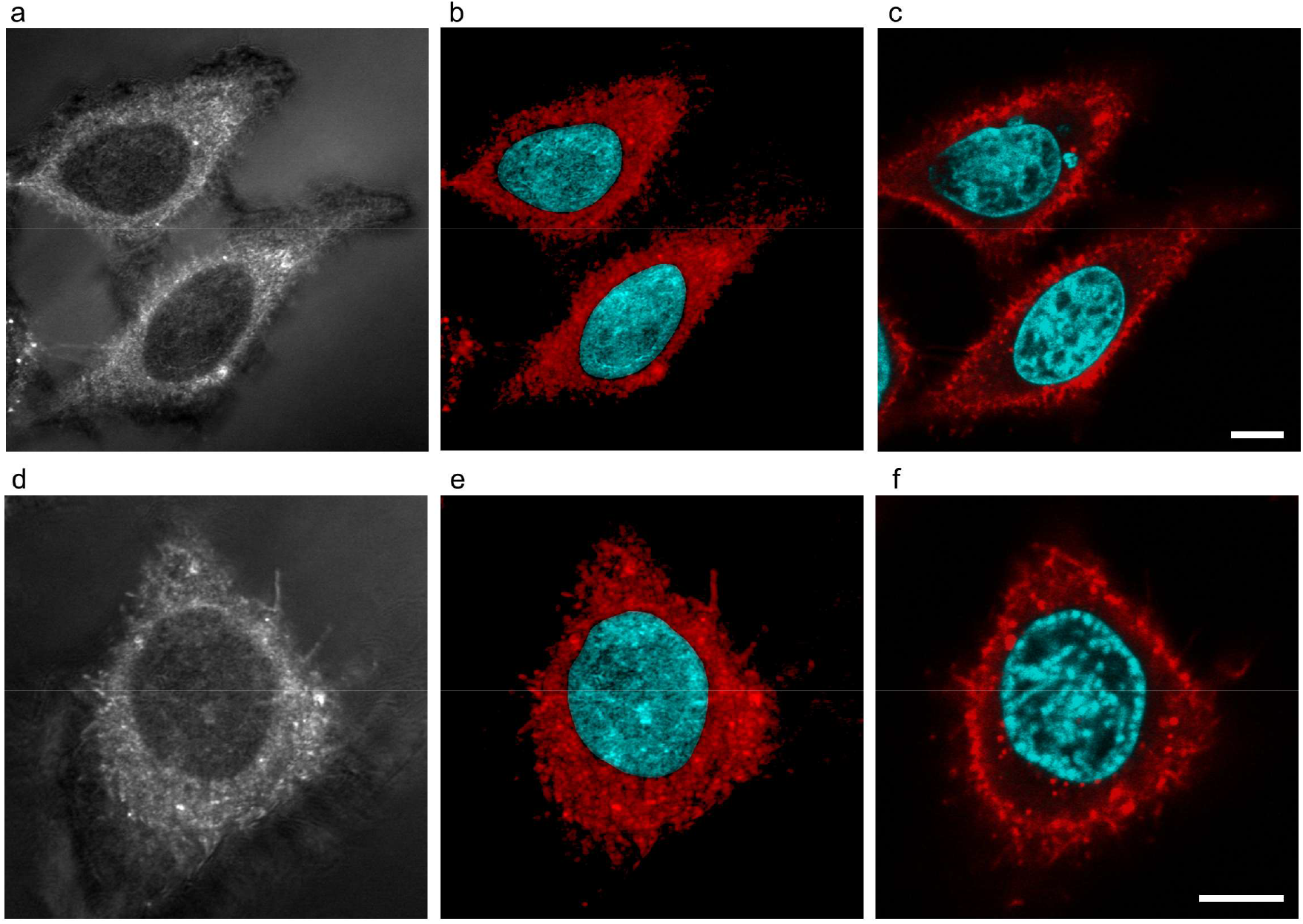
Pseudo-staining of label-free BBCM images. **a**, BBCM image of two HeLa cells showing clear cell and nuclear outlines. **b**, Pseudo-stained rendering of (a), mapping entropy-enhanced regions to red and nucleus-masked regions to cyan. **c**, Corresponding two-channel fluorescence image of the same field of view as (a), with DAPI-stained nuclei (cyan) and Concanavalin-A Alexa Fluor 647-labeled glycoconjugates on intracellular membranes and the plasma membrane (red). **d**, BBCM image of a HeLa cell. **e**, Pseudo-stained rendering of (d). **f**, Corresponding two-channel fluorescence image of the same field of view as (d), again with DAPI (cyan) and Concanavalin A-Alexa Fluor 647 (red). Scale bar 10 µm.

These results, validated across four cell lines (pancreatic PANC-1 cells: Fig. 2 and Supplementary Fig. 2, cervical HeLa cells: Fig. 3, Fig. 4, and Supplementary Fig. 3, ovarian SK-OV-3 cells: Fig. 5 and Fig. 6, and esophageal OE19 cells: Supplementary Fig. 2), demonstrate that BBCM reliably visualizes both cellular and nuclear outlines. Moreover, the images can be rendered in a familiar format that closely mimics conventional nuclear and cytoplasmic stains. Notably, acquiring the fluorescence images required no modifications to the microscope, highlighting the strong multimodal potential of BBCM. Our data indicates that fluorescence labelling does not interfere with BBCM performance across the tested cell types, and suggests that these findings are likely generalizable to other cell lines.

**Fig. 4.**
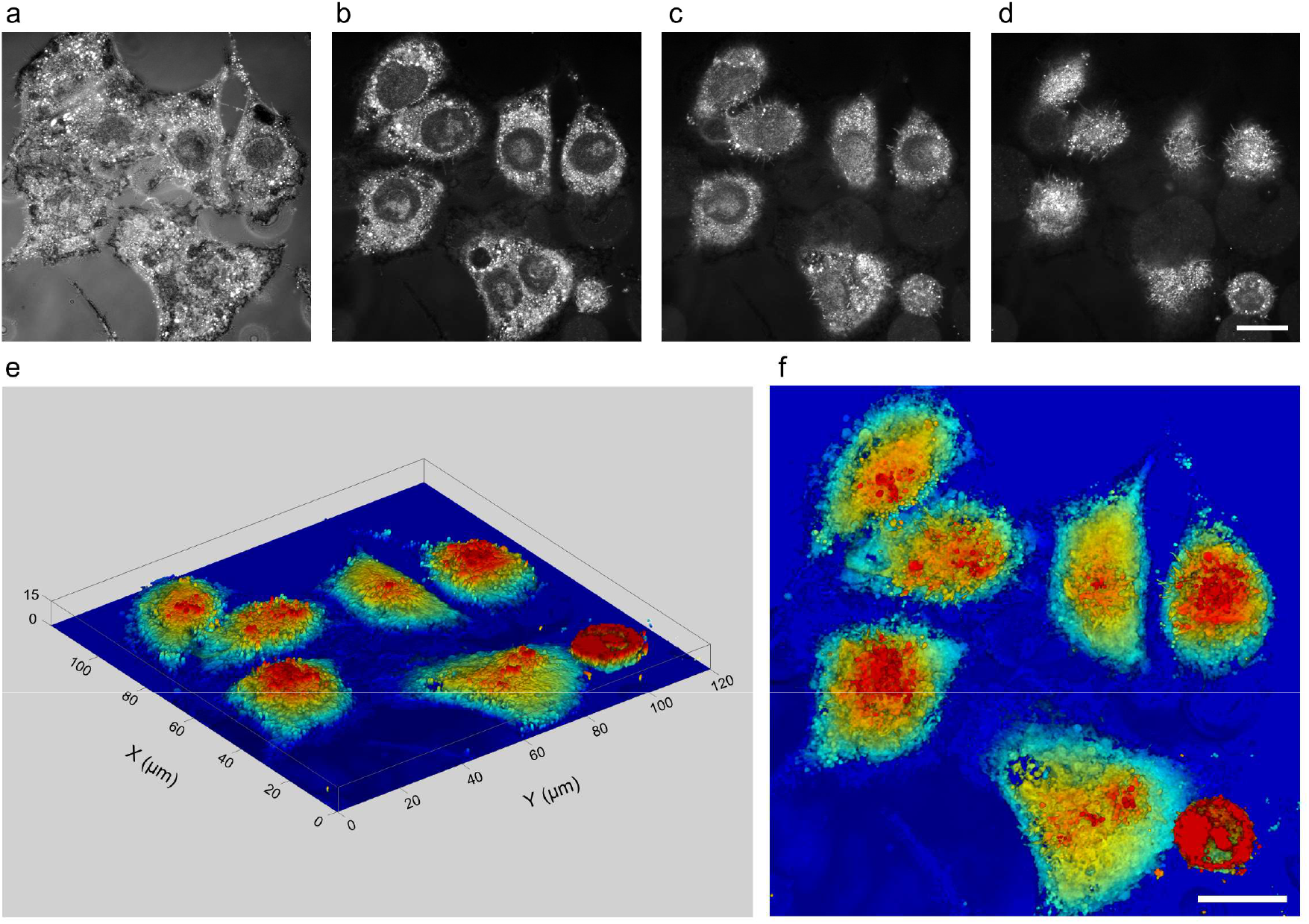
Label-free 3D cell reconstructions. **a**, A BBCM z-stack slice of HeLa cells at 1 µm above the glass surface, showing cell edges adherent to the surface. **b**, BBCM slice at 4 µm above the glass surface, showing clearly defined nuclear outlines. One cell appears to have been fixed during division. **c**, BBCM slice at 7 µm above the glass surface, near the upper region of the nuclei. **d**, BBCM slice at 10 µm above the glass surface, approaching the top of the cell. **e**, 3D reconstruction with depth encoding by color. **f**, Top-down view of the 3D reconstruction in (e). Scale bar 20 µm.

**Fig. 5.**
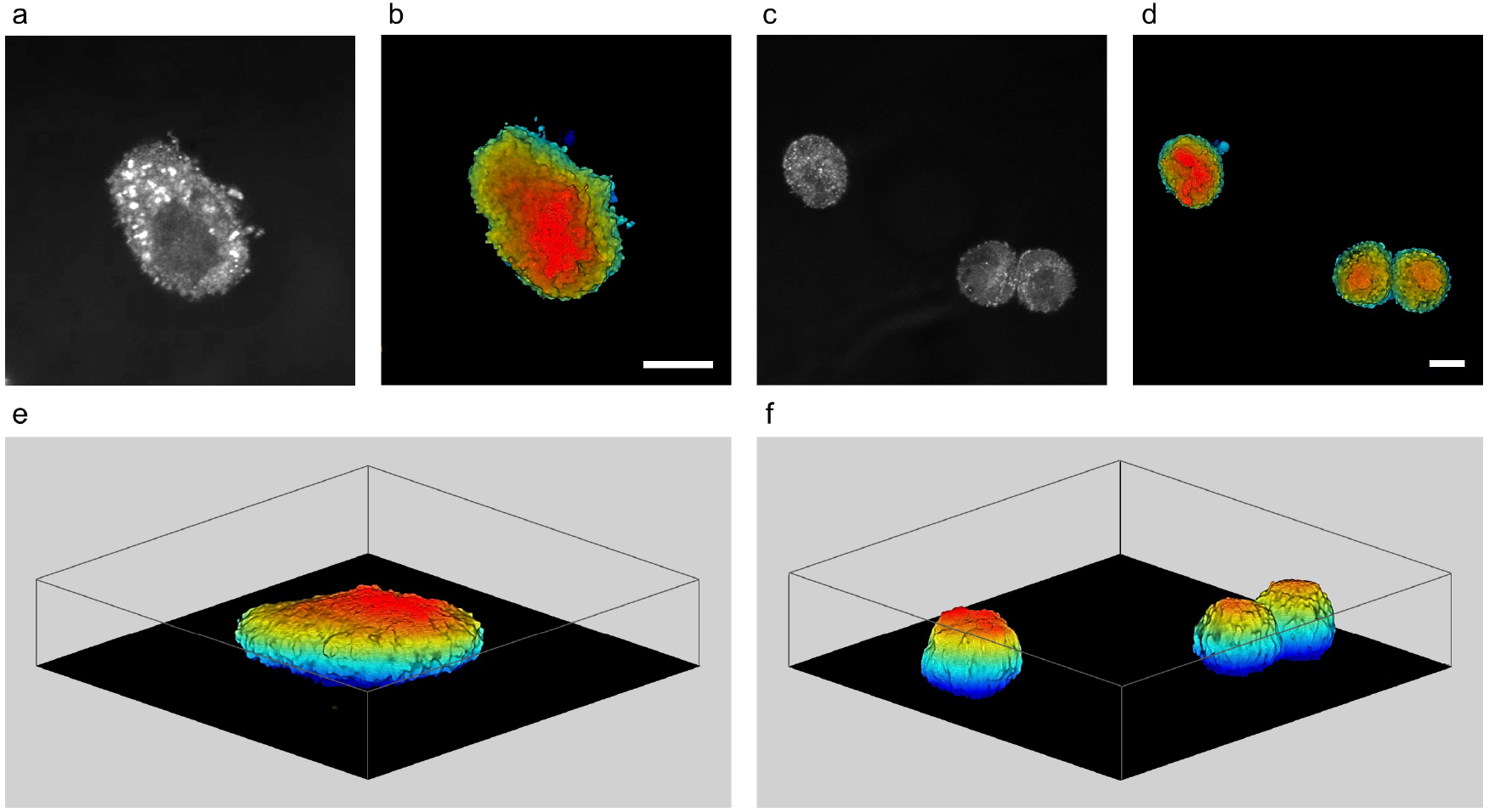
Label-free 3D reconstructions of cell suspensions. **a**, BBCM z-stack slice of a single SK-OV-3 cell. **b**, Same cell as (a) shown in a top-down view with depth encoded by color. **c**, Single z-slice of two SK-OV-3 cells, with one cell in an advanced stage of division. **d**, Same cells as (c) shown in a top-down view with depth encoded by color. **e**, 3D reconstruction of the cell shown in (a). Total z-height 23 µm. **f**, 3D reconstruction of the cell shown in (c). Total z-height 26 µm. Scale bars 10 µm.

**Fig. 6.**
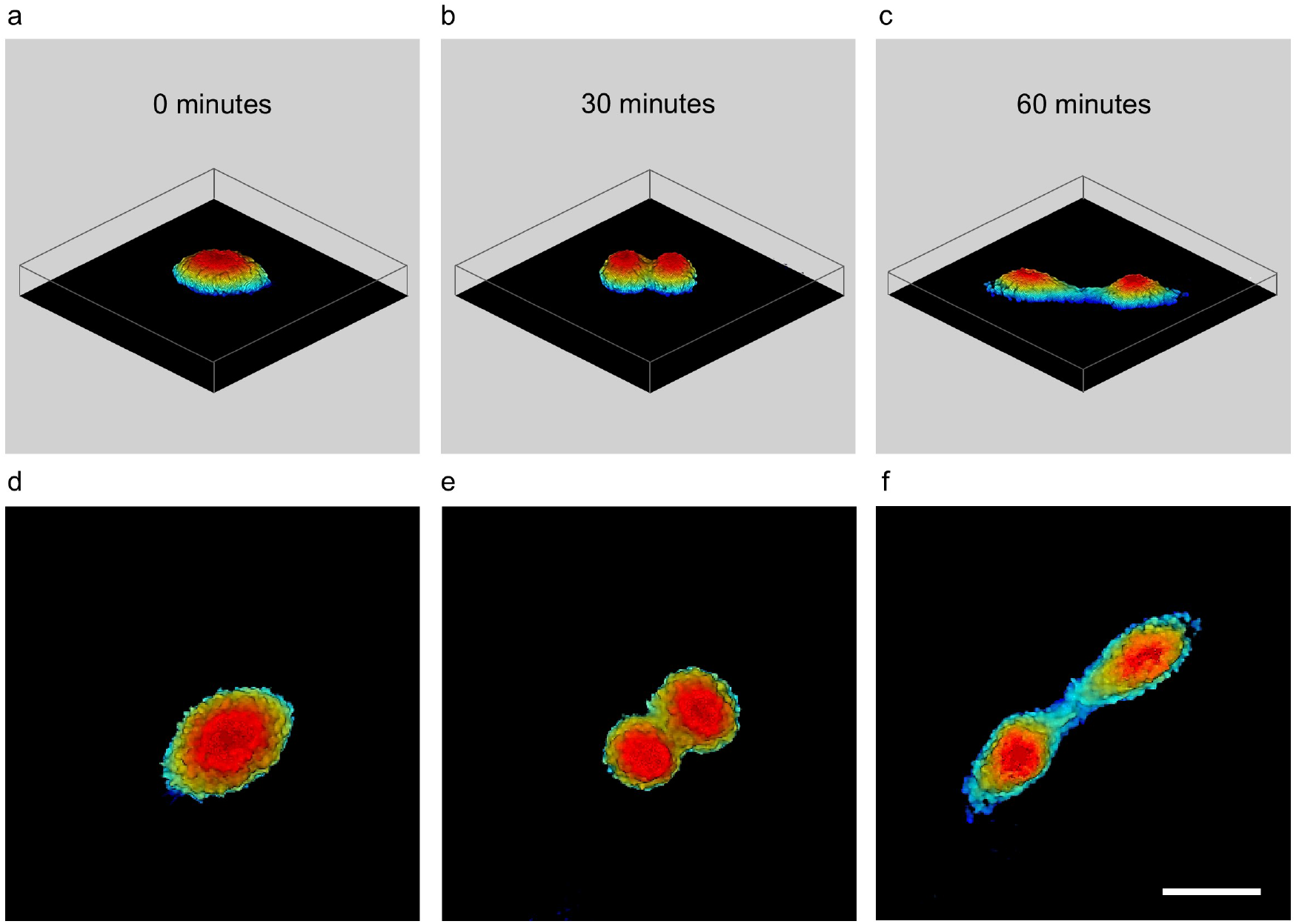
BBCM live-cell time-lapse imaging. **a**, 3D reconstruction of a single SK-OV-3 cell at t=0 minutes. Total z-height 21 µm. **b**, 3D reconstruction at t=30 minutes with the onset of division evident. Total z-height 20 µm. **c**, 3D reconstruction at t=60 minutes with two separate cells visible. Total z-height 15 µm. **d-f**, Top-down views with depth encoded by color at t=0 minutes, 30 minutes, and 60 minutes, respectively. Scale bar 20 µm.

### Label-free three-dimensional cell reconstructions

Given the optical sectioning provided by the confocal configuration, it is possible to perform label-free 3D reconstructions of cellular architecture, as shown in Fig. 4. A standard z-stack was obtained from a HeLa cell sample, with 31 slices acquired at a 0.5 µm spacing, resulting in a total z-height of 15 µm. Figures 4a-d show slices at 1 µm, 4 µm, 7 µm, and 10 µm above the glass surface, respectively. At 4 µm depth (Fig. 4b), both cell boundaries and nuclear outlines are clearly visible. Interestingly, the nuclear morphology shows that five of the six cells contain a single nucleus, while the sixth cell appears to have been fixed just prior to division.

To generate a high-contrast 3D reconstruction, we developed a custom MATLAB pipeline. Briefly, background intensity was removed by selecting a user-defined region away from the cells and subtracting its average intensity from each pixel in a slice specific manner. This correction accounts for depth-dependent background intensity near the glass surface. This is evident in Figs. 4a–d, where background intensity is stronger near the glass surface and falls off rapidly with depth, consistent with confocal pinhole rejection. After normalization, a single intensity threshold was applied to isolate cellular features. Each z-slice was rendered as a semi-transparent surface, with opacity scaled to local intensity and color encoded by axial position. The resulting 3D reconstruction is shown in Fig. 4e, with a corresponding top-down view in Fig. 4f. More details on the reconstruction algorithm are provided in the *Methods* section.

Importantly, this method is not limited to adherent cells. To demonstrate its versatility, we imaged SK-OV-3 ovarian cancer cells in suspension with BBCM (Fig. 5). A representative mid-plane slice of a single cell reveals clear delineation of the cell boundary and nuclear envelope (Fig. 5a), highlighting BBCM’s ability to resolve structural features in suspension. The same 3D reconstruction procedure was used, with the first optical slice manually set to black. The top-down view with color encoded depth is shown in Fig. 5b. A separate BBCM slice from a different region is shown in Fig. 5c, with the corresponding top-down depth-encoded view in Fig. 5d. This field of view contains a single cell in the top-left corner and another in a late stage of division in the bottom-right. Volumetric reconstructions of the cells in Figs. 5a and 5c are shown in Figs. 5e and 5f, respectively.

### Three-dimensional live cell imaging

The label-free nature of BBCM makes it well suited for long-term, high-contrast imaging of live cells in their native state. Although BBCM requires no fluorescence labeling, careful control of illumination power remains essential for live-cell compatibility. To determine the minimum power required for high-quality imaging, we inserted a neutral-density filter into the optical path and systematically reduced the incident power. We found that incident power below 100 µW with a 1.6 µs pixel dwell time was sufficient to produce high-contrast label-free images with no detectable loss of performance. These illumination levels are well below established photodamage thresholds, indicating BBCM’s suitability for non-invasive, long-term imaging of live cells. To validate this, we cultured live SK-OV-3 cells and imaged them using an onstage incubation system integrated with the BBCM platform. Figure 6a shows a single cell at the start of the time course, and Fig. 6b shows the same cell 30 minutes later, exhibiting clear morphological hallmarks of mitosis. By

60 minutes (Fig. 6c), the cell has completed division, with both daughter cells distinctly resolved. Corresponding top-down, depth-encoded views of the same time points are shown in Figs. 6d-f, further highlighting BBCM’s ability to capture the dynamic 3D cellular environment without exogenous labels.

## DISCUSSION

BBCM exploits broadband light scattering to generate high-contrast label-free images. The technique employs a supercontinuum light source, which provides a bright, spatially coherent, well collimated broadband beam that can be focused into a small focal volume. The confocal pinhole rejects backscattered light from outside the confocal volume, with intensity determined by the number of scatterers present, as well as their shape, orientation, and refractive index. BBCM operates on principles similar to reflectance confocal microscopy (RCM), where endogenous scatterers such as melanin provide strong contrast and make cellular structures readily visible in vivo^33^. RCM has found widespread use in dermatology^34^ and has also been applied in experimental biology, for example to characterize three-dimensional collagen lattice reorganization during melanoma cell migration^35^. While RCM performs well in melanin-rich tissues or collagenous structures, its contrast depends on the presence of strong endogenous scatterers. In contrast, BBCM uses broadband illumination to spectrally average backscattered signal, enabling high-contrast, label-free imaging of diverse cell types and subcellular structures even in the absence of melanin or other highly scattering components. This spectral averaging suppresses the size-dependent oscillations that occur with monochromatic backscattering, removing wavelength-specific nulls and intensity overlaps. As a result, all scatterers produce measurable signal that increases approximately linearly with size, yielding high-contrast images even in cells with weak intrinsic contrast. This is illustrated in Supplementary Fig. 3, where we compare cells imaged with monochromatic and broadband illumination.

The optical sectioning provided by the confocal configuration in BBCM allows the internal structure of the cell to be easily probed. One important structure that can be analyzed is nuclear morphology, which is a central marker in cell biology. Furthermore, given that both cell and nuclear outlines can be easily observed, and because co-registered fluorescence images can be acquired, this technique has strong potential for deep learning-based virtual staining approaches^36-39^. The optical sectioning also allows straightforward three-dimensional reconstructions. While many label-free imaging techniques, such as dark-field, phase contrast, and DIC, can visualize live cells in two dimensions, they do not capture the full three-dimensional architecture of cells and tissues. Three-dimensional imaging is essential in contexts where spatial relationships and depth information fundamentally impact interpretation. For example, observing cell-cell interactions, such as T-cell engagement and killing of target cells^40,41^, requires volumetric imaging to confirm physical contact. Similarly, accurate measurement of cell volume, nuclear morphology, and membrane protrusions is often confounded in 2D projections. Though not shown here, BBCM is also applicable to 3D cell cultures like organoids and tumor spheroids^42,43^. In such structures, three-dimensional imaging enables clear segmentation and tracking of individual cells, which would otherwise be impossible with 2D techniques.

To create the three-dimensional reconstructions, we used standard z-stack acquisitions and a straightforward algorithm. The first step involved background subtraction, followed by a threshold to extract cellular detail. Other segmentation methods could also be easily applied here, for example more advanced model-based approaches^44^ or deep learning-based techniques^45^. After segmentation of the cellular detail in each z-stack slice, the images are stacked to reconstruct a three-dimensional volume. Here the z-height is obtained from the physical movement of the microscope objective, and does not require any assumptions on the refractive index of the specimen.

We have introduced a label-free technique for three-dimensional size-sensitive imaging of live cells. We validated its performance across diverse cell types, including adherent cells and cell suspensions. We have also shown that BBCM can be used at low optical powers, making it suitable for live cell time-lapse imaging. BBCM is simple to retrofit on existing confocal systems and is cost-effective, with the only additions being a standard beamsplitter and a supercontinuum light source, with entry-level sources available for less than $10,000. Because BBCM uses broadband illumination, it avoids the blind spots and ambiguities inherent to monochromatic backscattering, enabling robust size-sensitive contrast. BBCM integrates readily with multimodal measurements, is straightforward to use by nonspecialists, and opens new opportunities for quantitative, physiologically relevant live cell imaging.

## METHODS

### BBCM system setup

A Zeiss LSM 510 META laser scanning confocal microscope frame was used. The system utilizes a broadband coherent supercontinuum source (WhiteLase Micro, NKT Photonics). The visible (400 – 850 nm) output power of the supercontinuum source is 25 mW and the repetition rate is 30 MHz. A SCHOTT KG1 (Edmund Optics), 1 mm thick, colored glass heat absorbing shortpass filter was used to remove the infrared portion of the spectrum. The collimated visible light beam from the supercontinuum source was aligned with free-space coupling optics at the microscope entrance port and passes through a standard 80/20 beamsplitter. A high NA = 1.4 achromatic microscope objective (Zeiss Plan-Apochromat 63X) with low chromatic aberration was used to focus the beam. The spectrum after the objective was measured using an integration sphere and commercial spectrometer (AvaSpec), showing that light between 425 nm and 800 nm successfully travelled through the microscope. The backscattered light passes through a collection pinhole, which blocks most of the light originating above and below the focal plane. This allows only the light scattered within the focal volume to be detected. The white light point spread function of the microscope was measured with diameter 100 nm Polybead polystyrene microspheres (Polysciences), using standard methods^46^. The resulting data is shown in Supplementary Fig. 4, with the x, y, and z full width half maximum dimensions being 256 nm, 262 nm, and 1043 nm, respectively. The pixel dwell time for all cell measurements was 1.6 µs, which allows a 512 x 512 image to be acquired in less than 1 second.

### Microsphere measurements

Diameter 100 nm and 350 nm Polybead polystyrene microspheres (Polysciences) were used for the Mie scattering measurements. The microspheres (100 nm or 350 nm) were each mixed with 150 µL of 1% agarose at a 1:1000 ratio and dispensed onto the center of glass-bottom dishes (MatTek). The dishes were stored at 4 °C until the agarose solidified, forming an approximately 1 mm thick gel containing microspheres. The spectral measurements were acquired with the built in 32 channel PMT-based META spectrometer, which detects signals in the 411 nm to 754 nm spectral range with a 10.7 nm spectral resolution. MATLAB (MathWorks) code was written to extract the spectra from in-focus microspheres. The resulting spectra contained contributions from the microsphere properties (i.e. diameter and refractive index) along with contributions from the supercontinuum structure, which is shown in Fig. 1c. Dividing the average spectra from microspheres of different diameters removes the effect of the supercontinuum, given that the supercontinuum structure is constant across all measurements. This is illustrated in Supplementary Fig. 5, where Supplementary Fig. 5a shows the average raw spectra for diameter 100 nm and 350 nm microspheres. The open circles in Supplementary Fig. 5b show the divided spectrum, which is free from supercontinuum structure.

### Cell culture and fixation

HeLa cells were cultured in T-75 flasks (Thermo Fisher Scientific) using Dulbecco’s Modified Eagle Medium (DMEM) supplemented with 10% fetal bovine serum (FBS). Cultures were maintained at 37 °C in a humidified incubator with 5% CO_2_. The cells were plated onto µ-Slide 8 Well chambers (ibidi) at 30k cells per each well. After 24 hours, cells were fixed with 3% (w/v) paraformaldehyde (PFA) and 0.1% glutaraldehyde in phosphate buffered saline (PBS) for 10 minutes, quenched with freshly prepared 0.1% sodium borohydride in PBS for 7 minutes, and washed three times with PBS at room temperature. PANC-1 and OE19 cells were cultured in glass-bottom dishes (MatTek) using DMEM supplemented with 10% FBS. Cultures were maintained at 37 °C in a humidified incubator with 5% CO_2_. When cells reached approximately 50% confluency, they were fixed with 4% PFA for 8 minutes at room temperature, followed by three washes with PBS. SK-OV-3 cells were cultured in McCoy’s 5 A modified medium (Thermo Fisher Scientific) supplemented with 10% FBS. When cultures reached approximately 70% confluency, cells were detached by incubation with 1 mL trypsin–EDTA solution (Thermo Fisher Scientific) for 10 min or until the cell layer was dispersed. The enzymatic reaction was quenched by adding 6 mL of complete growth medium. Cells were collected by centrifugation at 800 rpm for 5 minutes, and the resulting pellet was resuspended in McCoys 5A without phenol red (Innovative Research) for subsequent use.

### Fluorescent labeling

HeLa and PANC-1 cells were processed for fluorescent labeling following fixation as described above. All staining steps were carried out at room temperature unless otherwise stated, and all solutions were prepared fresh on the day of use. Fixed samples were washed three times with PBS (pH 7.4; 5 minutes each) to remove residual fixative. For HeLa cells, the fixed cells were permeabilized in blocking buffer (3% bovine serum albumin (BSA), 0.1% TritonX-100 in PBS) for 10 minutes, followed by three washes in PBS. Cells were incubated with Concanavalin-A Alexa Flour 647 (Thermo Fisher Scientific, Cat. # C21421) at 100 µg/mL in PBS for 20 minutes under light-protected conditions with gentle rocking, then washed three times with PBS (5 minutes each). Nuclei were counterstained with DAPI (Sigma-Aldrich, Cat. # D9542) at 1 µg/mL in PBS for 5 minutes at room temperature on the rocker, followed by three additional PBS washes. For PANC-1 cells, nucleic acids were stained after fixation using DAPI (Sigma-Aldrich, Cat. # D9542) at a final concentration of 1 µg/mL in PBS for 5 minutes at room temperature. After incubation, the staining solution was removed, and cells were washed three times with PBS (5 minutes each). Staining was carried out under subdued lighting conditions to minimize fluorophore degradation. Each staining experiment was independently repeated at least three times to ensure reproducibility. Consistent staining patterns were observed across biological replicates for all cell lines. Samples were imaged in PBS (pH 7.4) at room temperature.

### Pseudo-stain code

To generate pseudo-stained images highlighting cell nuclei against the surrounding cellular background, we developed a custom MATLAB (Mathworks) script. BBCM images were first normalized and processed with a local entropy filter (window size = 9 pixels) to enhance fine structural detail. The resulting texture-enhanced image was multiplied with the original BBCM image, providing an image with a uniform low-intensity background. Pixel intensities were mapped to a red color channel using an offset and gamma correction, while the green and blue channels were set to zero. The resulting images are provided in Supplementary Fig. 6, showing that the above steps generate a pseudo-stained image similar to what is observed for cytoplasm-stained fluorescence images. Next, a binary nucleus mask, obtained from prior segmentation, was used to separate nuclear and non-nuclear regions. Intensities from the original BBCM image within the mask regions were extracted, normalized to their local dynamic range, adjusted with gamma correction, and mapped equally to the green and blue channels. Finally, to avoid sharp edges between nuclear and background regions, a distance transform was applied to the nucleus mask, allowing a gradual transition from cyan to red at the nuclear boundary.

### 3D reconstruction code

To generate high contrast 3D reconstructions from raw z-stacks, we developed MATLAB (MathWorks) code that was optimized for BBCM. The code first imports the LSM z-stack and crops it to a user-defined subset of slices. This allows exclusion of planes above the cells or planes below the glass surface. To address slice-to-slice background variation, a background region is selected manually on the first slice, and its average pixel intensity is used to subtract slice-specific background levels across the entire stack. Each slice is then clipped to remove negative values and normalized to the global maximum intensity. To enhance continuity, we applied a 3×3×3 box filter and interpolated along the z-axis. For visualization, each slice is converted into a semi-transparent surface, where the color encodes depth and the opacity is determined by the relative intensity of each voxel. Finally, to prevent high-intensity outliers from dominating the rendering, we scale the opacity using a percentile-based saturation scheme.

### Live cell imaging

The laser power was reduced using a neutral density filter (NE20B-A, Thorlabs). To measure the optical power, the objective was removed and the collimated beam was directed into an optical power meter (PM100D with S121C sensor, Thorlabs). After the laser stabilized, the resulting power at a 600 nm setting was 89 ± 3 µW. Given the responsivity of the sensor, and red-weighted spectral shape, this is expected to be an over-estimate of the optical power. Along with reduced optical power, an on-stage incubator (Okolab) was used to maintain the cell viability. This system included a temperature controller, gas controller, incubating chamber, humidity module, and objective heater. SK-OV-3 cells were cultured in glass bottom dishes (MatTek) and when the cells reached approximately 50% confluency, the medium was replaced with complete growth medium without phenol red. The dishes were then transferred to the on-stage incubator for BBCM measurements. Z-stacks were acquired every 30 minutes over a 12-hour period.

## Data availability

The main data supporting the results of this study are available within the paper. Raw z-stacks, along with source data for all plots, are available from the corresponding authors upon reasonable request.

## Code availability

All imaging data was acquired using standard Zeiss imaging software, with no custom acquisition code required. Code for generating pseudo-stained images and 3D reconstructions is available via GitHub at https://github.com/mfcoughl/BBCM.

## ACKNOWLEDGEMENTS

This work was supported by US National Institutes of Health grants R01 EB025173, R01 CA228029, R01 CA293050, and R21 AG085089 and US National Science Foundation grants CBET 2220273, and CBET 2325317.

## AUTHOR CONTRIBUTIONS

L.T.P. and M.F.C. initiated and supervised the project; M.F.C., L.T.P. and L.Z. conceived the method; M.F.C, L.Q., Y.Z., P.K.U, X.Z and U.K constructed the imaging system; L.Z. and R.T.P. cultured cells, performed staining, and prepared samples for imaging; M.F.C and L.Z. performed imaging experiments; M.F.C, L.Z and L.T.P performed the data analysis; M.F.C developed algorithms and software; P.K.U., X.Z., R.T.P, L.Q. and L.T.P. evaluated the method; U.K. performed graphic design; M.F.C and L.T.P. wrote the manuscript with input from R.T.P and L.Z.

