## Supplementary Information for "Broadband backscattering confocal microscopy enables label-free 3D live cell nanoscale sensitive imaging"

[2]Harvard Biophysics Program
Harvard University

### SUPPLEMENTARY INFORMATION

[†] These authors contributed equally

### Contents

Supplementary Figures ........................................................................ 3

Microsphere Scattering Model ................................................................... 9

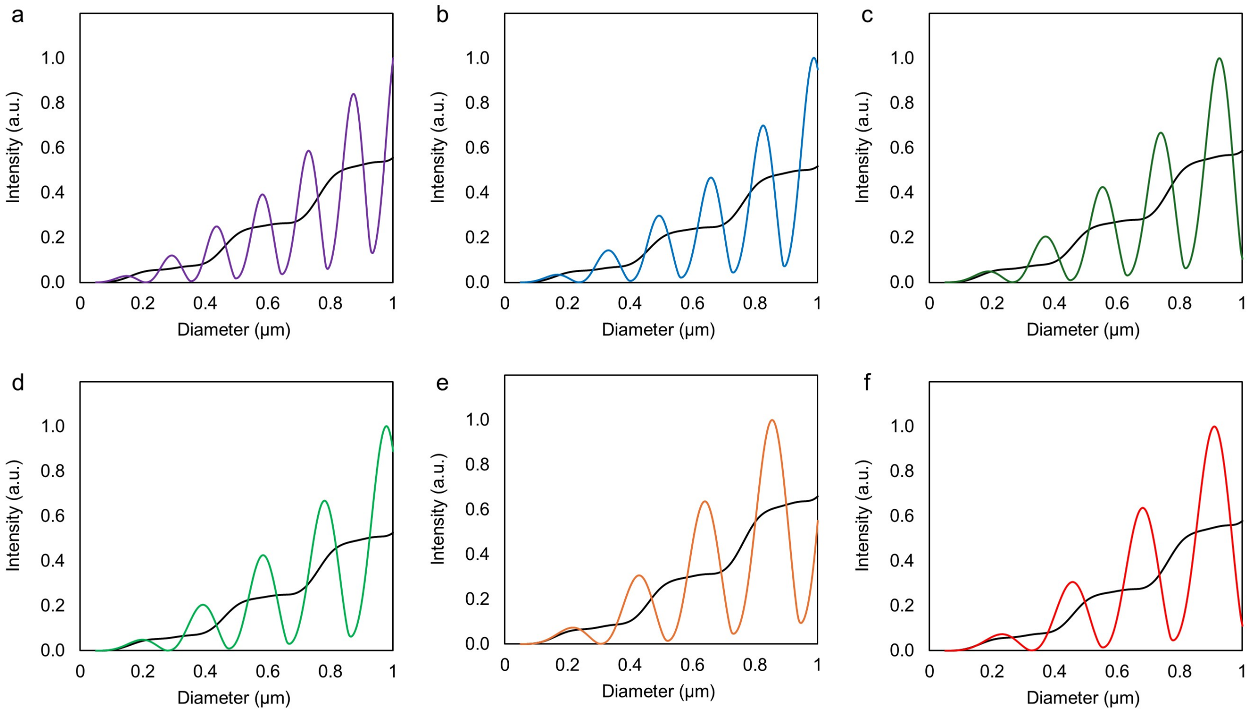


**Supplementary Fig. 1. Backscattering intensity vs particle size for various wavelengths. a,** Simulated backscattering intensity for spherical particles of varying diameters. The purple curve corresponds to 405 nm, while the black curve corresponds to broadband illumination in the 425 nm to 800 nm range. **b,** Monochromatic light at 458 nm. **c,** Monochromatic light at 514 nm. **d,** Monochromatic light at 543 nm. **e,** Monochromatic light at 594 nm. **f,** Monochromatic light at 633 nm.

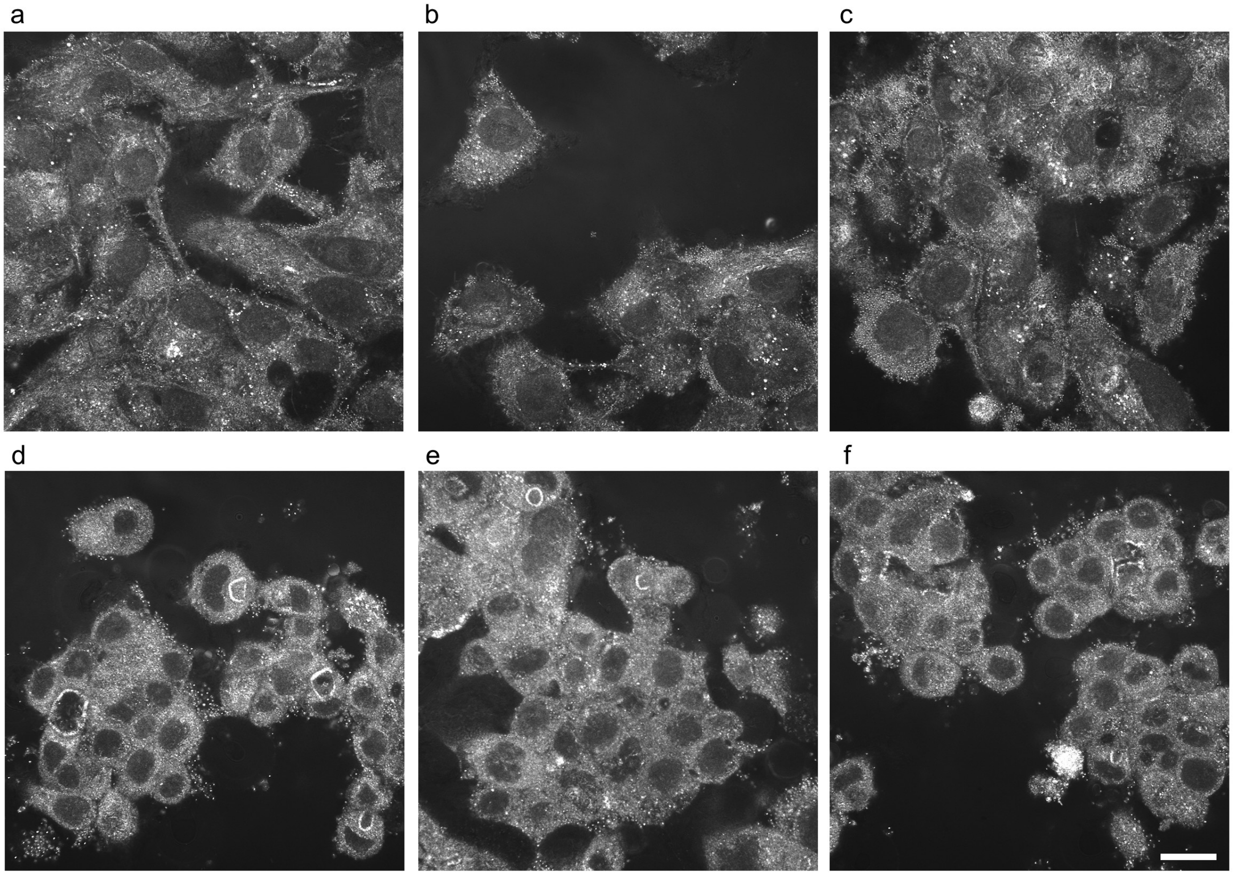


**Supplementary Fig. 2. Label-free BBCM images of unstained PANC-1 and OE19 cells. a-c,** BBCM images of PANC-1 cells, showing clear cell and nuclei outlines. **d-f,** BBCM images of OE19 cells, which again show clear cell and nuclei outlines. Scale bar 20 μm.

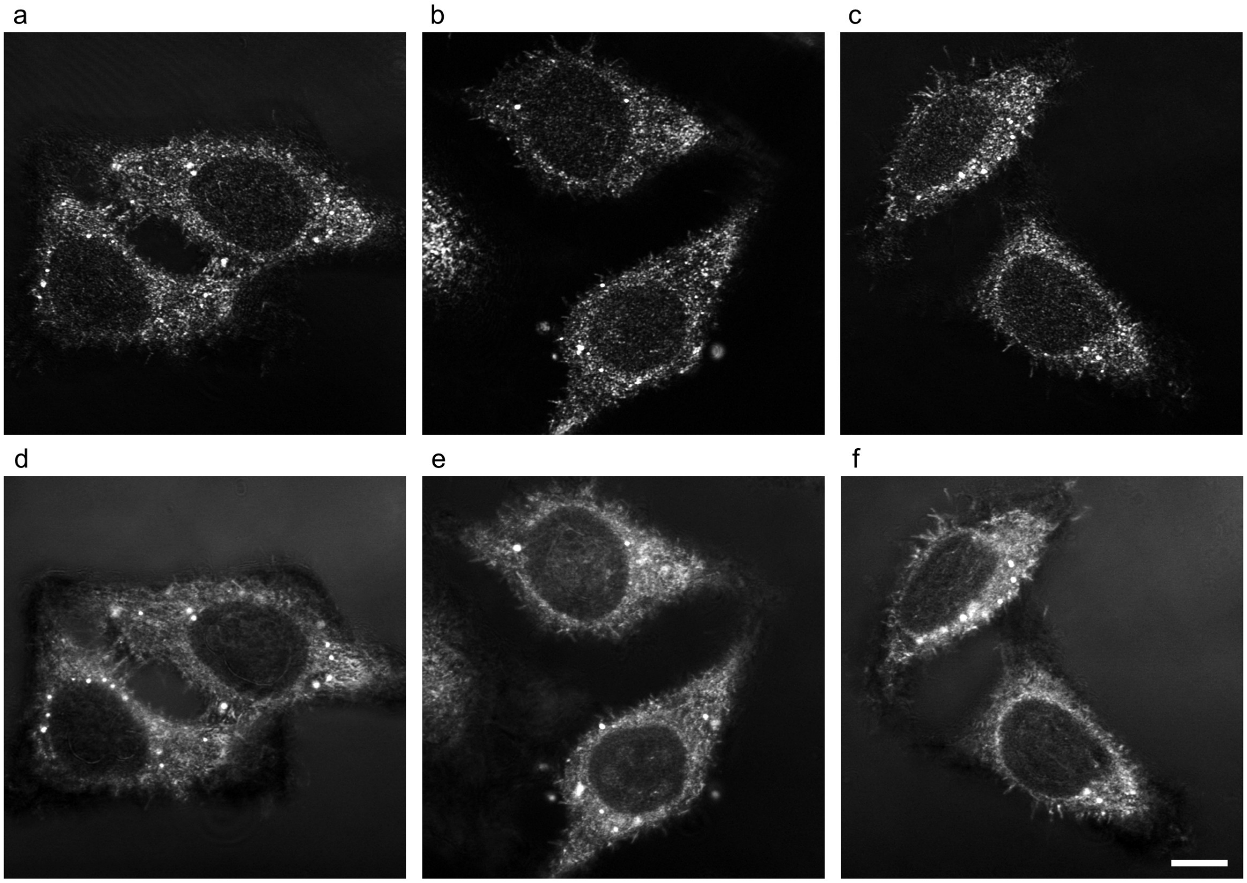


**Supplementary Fig. 3. Monochromatic and broadband illumination comparison. a-c,** Images of HeLa cells acquired with 488 nm illumination and processed in the same way as BBCM images. **d-f**, Corresponding BBCM images. Scale bar 10 µm.

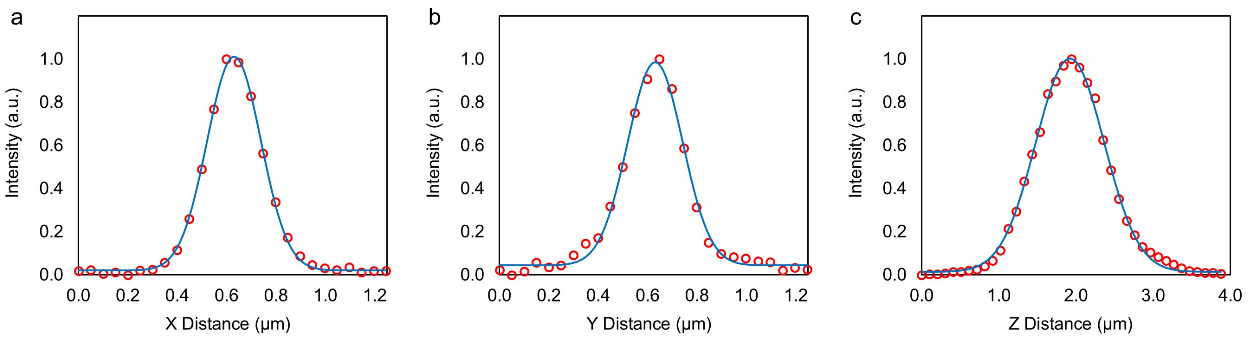


**Supplementary Fig. 4. White-light point spread function (PSF) measurements. a,** PSF measured with 100 nm-diameter polystyrene microspheres. The full width at half maximum (FWHM) of the PSF in the X direction was 256 nm. **b,** The PSF FWHM in the Y direction was 262 nm. **c**, The PSF FWHM in the Z direction was 1043 nm.

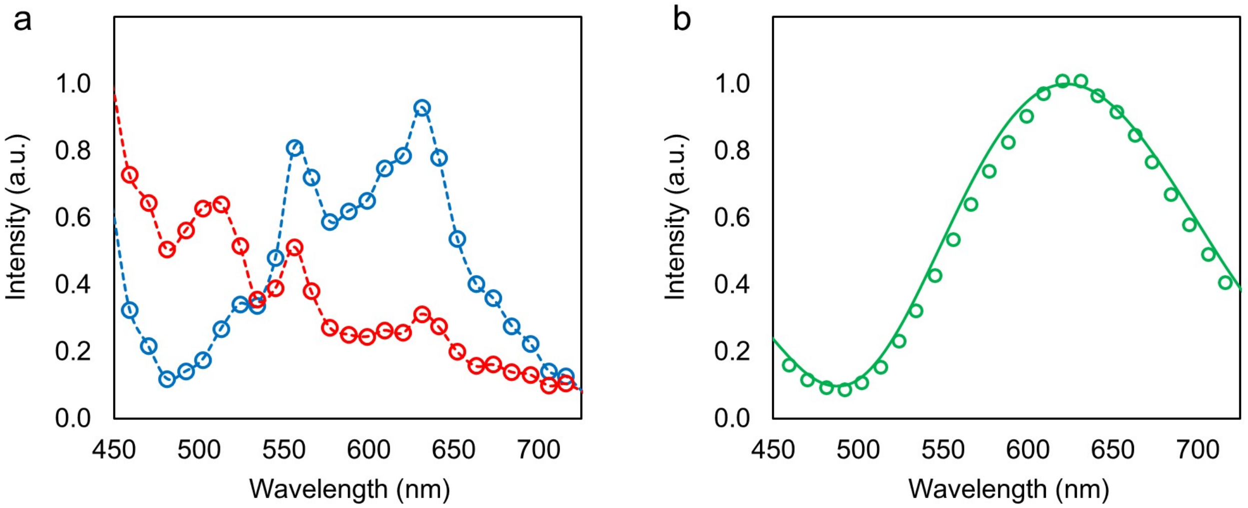


**Supplementary Fig. 5. Polystyrene microsphere measurements. a,** Average spectra for 350 nm-diameter polystyrene microspheres (blue circles) and 100 nm-diameter polystyrene microspheres (red circles). The dashed curves show the overall spectral shape for each microsphere size. The experimental spectra reflect a combination of the microspheres' scattering response and spectral features inherent to the supercontinuum light source. **b,** Dividing the 350 nm spectrum by the 100 nm spectrum removes the supercontinuum structure, which is constant across both measurements. The open circles represent the resulting normalized spectrum, while the solid curve shows the best fit using Mie theory. The Mie model incorporates angular averaging to account for the large numerical aperture of the objective.

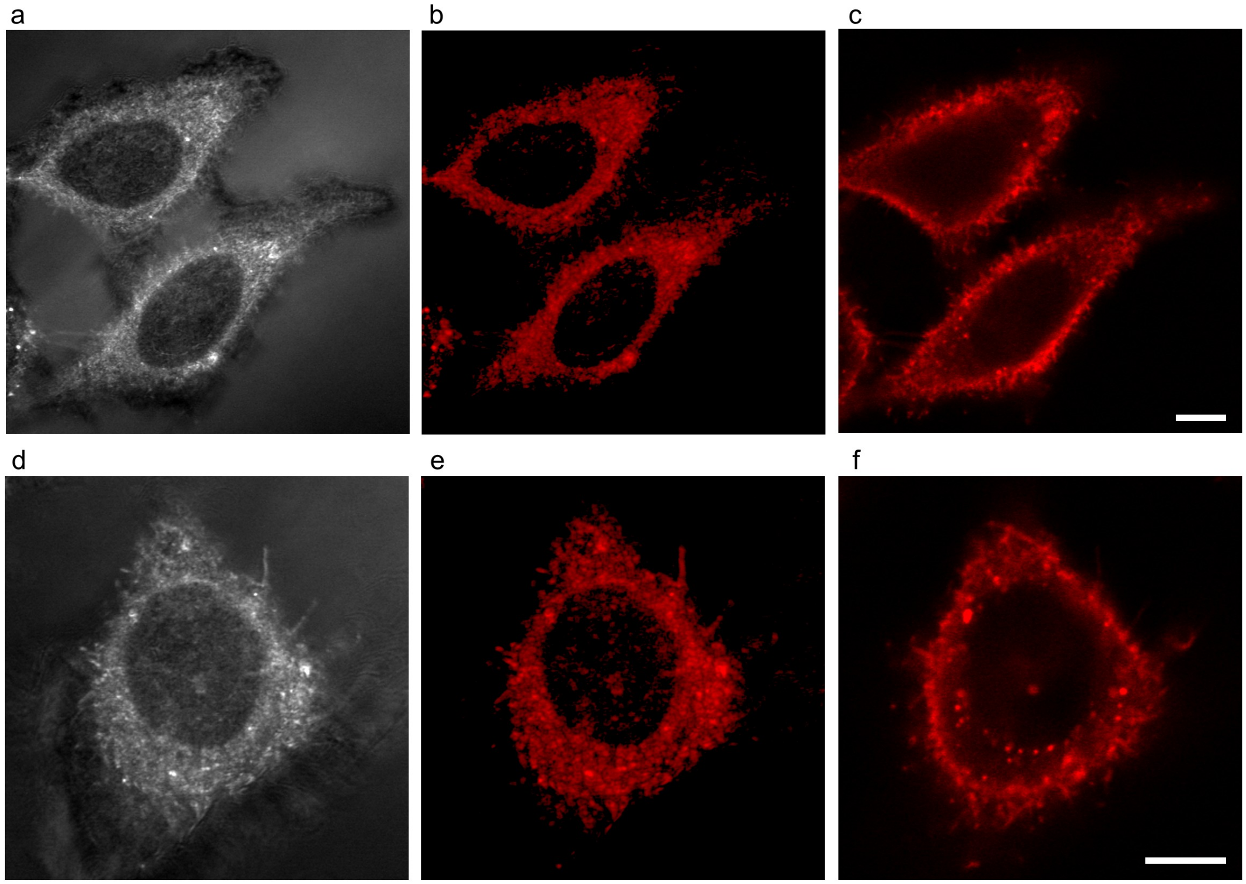


**Supplementary Fig. 6. Pseudo-staining of label-free BBCM images. a,** BBCM image of two HeLa cells showing clear cell and nuclear outlines. **b,** Pseudo-stained image of (a) mapping entropy-enhanced regions to red. **c,** Corresponding fluorescence image from the same field of view as (a), with Alexa Fluor 647–conjugated Concanavalin A labelling glycoconjugates on intracellular membranes and the plasma membrane (red). **d,** BBCM image of a single HeLa cell. **e,** Pseudo-stained image of (d). **f,** Fluorescence image from the same field of view as (d). Scale bar 10 μm.

#### Microsphere scattering model

The scattering model used here was published previously[28] and is not part of the BBCM method. Instead, it is used to illustrate BBCM's operating principle and to simulate how the BBCM signal intensity depends on particle size across wavelengths. The model accounts for the illumination and backscattering angle ranges set by the objective's numerical aperture and the wavelength dependence of coherent scattering. For polystyrene microspheres in water, Mie scattering theory[47] is used because polystyrene and water have a large refractive-index mismatch. In contrast, for subcellular compartments within the cytoplasm, where the refractive-index mismatch is much smaller, the scattering model can be simplified using the Rayleigh–Gans approximation[48].

To simulate the coherent scattering behavior of polystyrene microspheres embedded in agarose, we developed a MATLAB (MathWorks) script that models wavelength-dependent backscattering intensity under realistic optical conditions. The model assumes coherent illumination, in which incident plane waves arrive over a defined angular range, and the resulting electric fields are summed coherently before computing the total backscattered intensity. Backscattered fields were computed using MATLAB functions by Mätzler[49], which implement Mie theory for homogeneous spheres and provide robust estimates of scattering amplitudes and internal fields. For each wavelength and illumination angle, we evaluated the field contributions across the full backscattering hemisphere and averaged over all angles. A least-squares fit was used to match the simulated results to the experimental data in Supplementary Fig. 5b. The primary fit parameters were the microsphere diameter (considering that the manufacturer provides an average diameter ± standard deviation) and the angular range. The angular range could not be calculated directly from the objective's numerical aperture because of the beam's Gaussian profile and was therefore estimated by fitting. After finding the best fit for the data in Supplementary Fig. 5b, the returned angular range and microsphere diameters were used to generate the plot in Fig. 1d.

To simulate the backscattering intensity for microspheres with diameters between 50 nm and 1000 nm, the same code discussed above was used. However, the refractive indices were adjusted to more closely match biological conditions, with the microsphere refractive index set to 1.42 and the surrounding medium to 1.38. A one-dimensional wavelength array, corresponding to the microscope's transmitted spectral range, was used to calculate the backscattering spectrum for a given microsphere diameter. The intensity was then averaged across wavelengths, enabling straightforward comparison between broadband and monochromatic backscattering conditions. Backscattering intensity was calculated for the entire range of microsphere diameters, and the resulting intensities were normalized to the maximum. The final plots are shown in Fig. 1e and Supplementary Fig. 1.
